# The proposed Baltic Sea endemic green alga *Monostroma balticum* represents a foliose morphotype of *Ulva intestinalis* (Ulvales, Ulvophyceae)

**DOI:** 10.64898/2026.09.16.751975

**Authors:** Niko R. Johansson, Jaanika Blomster, Tytti Wärri, Elina Laiho, Elena Schrofner-Brunner, Nicolas Straube, Cessa Rauch, Sophie Steinhagen

**Author notes:** senior author (Sophie Steinhagen).

## Abstract

Green algae in *Ulva, Monostroma* and similar genera are notoriously difficult to identify morphologically, due to high phenotypic plasticity and the presence of cryptic species – which has historically resulted in a challenging taxonomy, poorly resolved species boundaries, and potentially numerous invalid species descriptions. DNA-based specimen identification, especially combined with analysis of historical herbarium voucher DNA, has huge potential in resolving many unresolved taxonomical problems in these algae. One such example is *Monostroma balticum,* which has been described as foliose, monostromatic green seaweed known only free-floating, and reported to be endemic to the Baltic Sea. Previous studies have highlighted major uncertainty in the status of this taxon. We conducted an extensive sampling campaign of monostromatic green algae across the Baltic Sea to molecularly, morphologically and ecologically characterise this enigmatic taxon. Additionally, we investigated original exciccate material of *M. balticum* by using a museomic approach. We show that *Monostroma balticum* represents a monostromatic, foliose morphotype of the cosmopolitan and ubiquitous species *Ulva intestinalis.* This foliose growth form is seen in low salinity conditions, and, so far, reported only from the inner basin of the Baltic Sea. Our work shows that using morphology- and DNA-based museomic approaches for contemporary and historical specimens is a powerful combination to resolve long-standing taxonomical uncertainties and to produce new biodiversity information, even in relatively well-studied regions.

**Highlights:**

- *Ulva intestinalis* can grow in a monostromatic foliose morphotype.
- Foliose *Ulva intestinalis* appears linked to low salinity conditions.
- DNA-analysis of historical specimens helps solve taxonomical problems.

## INTRODUCTION

Green algae in the genus *Ulva* Linnaeus (Ulvophycae, Ulvales) (Linnaeus 1753) as well as morphologically similar taxa, e.g. *Monostroma* Thuret (Monostromataceae, Ulotrichales) (Thuret 1854) are notoriously difficult to identify and classify based on morphological features alone (Steinhagen et al. 2019b; Bartolo et al. 2022, Lagourgue et al. 2022, Tran et al. 2022). This is partly due to extreme phenotypic plasticity, dependent on environmental conditions, such as salinity, temperature, light or nutrient conditions (Tan et al. 1999, Gao et al. 2016). Additionally, many species are cryptic or semi-cryptic in nature, with taxa having very similar morphological features, but clearly different evolutionary origins (Hofmann et al. 2010, Steinhagen et al. 2019b, Bartolo et al. 2022). The taxonomy and nomenclature of these algae is based on decades of macro- and micromorphological studies, accompanied with life cycle experiments (Monotilla et al. 2018, Steinhagen et al. 2019a) and ultrastructural studies (Stewart et al. 1973). It is therefore not surprising, that recent DNA- and genomic studies have caused considerable change in the taxonomy and systematics of this group (Tran et al. 2022), given its morphologically cryptic nature. DNA-based analysis of type material has revealed long-lasting misapplication of commonly used names (Hughey et al. 2019, 2021, 2024, 2026; Hughey and Gabrielson 2022; van der Loos et al. 2025). Despite these efforts, there is still plenty of taxonomical and nomenclatural uncertainty left in several taxa in this group – especially, if specimen identification is based solely on morphology.

Many species of *Ulva* and *Monostroma* are ubiquitous in aquatic systems, and gained considerable interest from not only systematists, but also in the fields of bioeconomy, aquaculture and biopharmaceutical production (Hofmann et al. 2025, Toth et al. 2025). Occasionally, these taxa cause green tides (Wan et al. 2017, Lagourgue et al. 2022), which may become more prevalent with global environmental change (Gao et al. 2017, Feng et al. 2026). Hence, detailed and explicit species identification and nomenclature in these algae is of high importance.

*Monostroma* (Monostromataceae, Ulotrichales) is a widely distributed genus of green seaweeds, characterized by a strictly monostromatic, i.e. single-cell-layered thallus. Worldwide 30 species are recognised in AlgaeBase (Guiry & Guiry 2026), of which two are known from the Baltic Sea: the cosmopolitan, attached, early season species *Monostroma grevillei* (Thuret) Wittrock and the proposed endemic, free-floating, late season species *Monostroma balticum* Wittrock. Despite apparently clear morphological and phenological differences between these taxa, the taxonomy and species status of M. balticum are unclear – even given its assumed endemic status and related potential conservation concern. Originally recognized by Johan Erhard Areschoug as “*Ulva baltica*”, the taxon was described and published by Veit Brecher Wittrock based on material from the Slite port in Gotland in his 1866 monograph dissertation of *Monostroma* (Wittrock 1866). Wittrock attributes the name *Monostroma balticum* to Areschoug’s *Ulva baltica*, even if it was not validly published. However, as Guiry & Guiry (2026) pointed out, Wittrock’s description can be considered valid if the name is solely attributed to him, as *Monostroma balticum* Wittrock. Wittrock did not to designate a single holotype specimen but refers to a gathering in the exsiccate series *Algae Scandinavicae Exsiccatae Serie novae* (1861–1872), collected by T. O. Krok.

*Monostroma balticum* is characterised as a monostromatic, free-floating blade up to 15 cm across, exclusively seen in the Baltic Sea in typically sheltered locations in late summer, sometimes forming mass accumulations. The microscopic morphology of the blade is described to consist of tightly packed cells with 5–7 corners and a plate-like chloroplast clearly visible as a distinct band in cross-section of the blade (Wittrock 1866). Many macroalgal floras from the Baltic Sea region recognise and treat this species after Wittrock’s description, generally agreeing with his original description (Svedelius 1901, Levring 1940, Wærn 1952, Ravanko 1968, Tolstoy & Österlund 2003). Later workers report the species growing considerably larger (up to 1 m, Joakim Hansen pers. comm.) and the description of microscopical features is in general vaguer than in Wittrock’s original treatment. Since originally known only in the free-floating stage, Wittrock hypothesises that attached, juvenile individuals (akin to juvenile *Monostroma grevillei*) could be found (Wittrock 1866). However, up to date, no workers claim to have observed attached and/or juvenile thalli of *M. balticum*, apart from the work of Orvokki Ravanko from Finland, where she claims they correspond to the ontogeny of *M. grevillei* (Ravanko 1969).

Ample taxonomic confusion exists around *Monostroma balticum* in the Baltic Sea. While some workers consider it a separate species, some think of it as a free-floating or detached form *or M. grevillei*, or some species of foliose or free-floating *Ulva*. Previous work has demonstrated that green tides in the Baltic Sea can be caused by a monostromatic aberrant growth form of *Ulva intestinalis* Linnaeus, but the link of these findings to the taxon *M. balticum* has been speculative (Bäck et al. 2000, Blomster et al. 2002). Free-floating foliose green macroalgae are relatively rare in the inner Baltic Sea, so the growth form, ecology and phenology alone have been the main basis of identification for this taxon. This gives room for misidentification. No molecular data exists for this taxon from contemporary or historical material. Despite this taxonomic confusion, the taxon is reasonably common and generally recognised by practising phycologists and marine biologists in the region.

In this study, we aim to shed light to this proposed endemic taxon by characterising both original and contemporary material using an integrated approach of DNA-based methods, including museomics, morphology and ecology. To achieve this, we conducted a wide sampling campaign of foliose, monostromatic green algae across the Baltic Sea region to molecularly and morphologically characterise all corresponding taxa. By critically reassessing the original description of *Monostroma balticum* and examining historical herbarium material, we aim to clarify the taxonomic status of this putatively endemic species.

## MATERIALS AND METHODS

### Sampling

Foliose monostromatic macrophytes (= macromorphologically resembling *Monostroma* sensu lato) were collected across the Atlantic-Baltic Sea region including collection locations in four countries (Denmark, Finland, Norway, Sweden), spanning multiple ecosystems (Fig 1, Table S1). The sampling was designed to fully cover the known distribution of *M. balticum* (Wittrock 1866, Levring 1940, Wærn 1952, Tolstoy & Österlund 2003) across a wide geographical, phenological and ecological range. Additionally, sampling encompassed the broad salinity gradient characteristic of the Baltic Sea, ranging from marine conditions in areas connected to the North Atlantic to nearly freshwater conditions in the innermost parts of the Baltic Sea (Lehmann et al. 2022). Per sampling site, all monostromatic distinct algal forms were collected and photographed. A 1–2 cm2 tissue sample, clean from epiphytes, was collected in silica gel for genomic DNA extraction. Selected specimens were studied microscopically and preserved as herbarium vouchers. All herbarium specimens are deposited in relevant herbaria (see Table S1). Prevailing abiotic parameters (salinity, temperature, oxygen concentration) during collection were documented from sampling sites in Sweden, Denmark and Norway and were retrieved from the Baltic Sea Physics Reanalysis dataset from the EU Copernicus Marine Service Information database (CMEMS 2026) for other samples. In addition, relevant samples from van der Loos et al. 2022 were included in our treatment.

**Fig 1:**
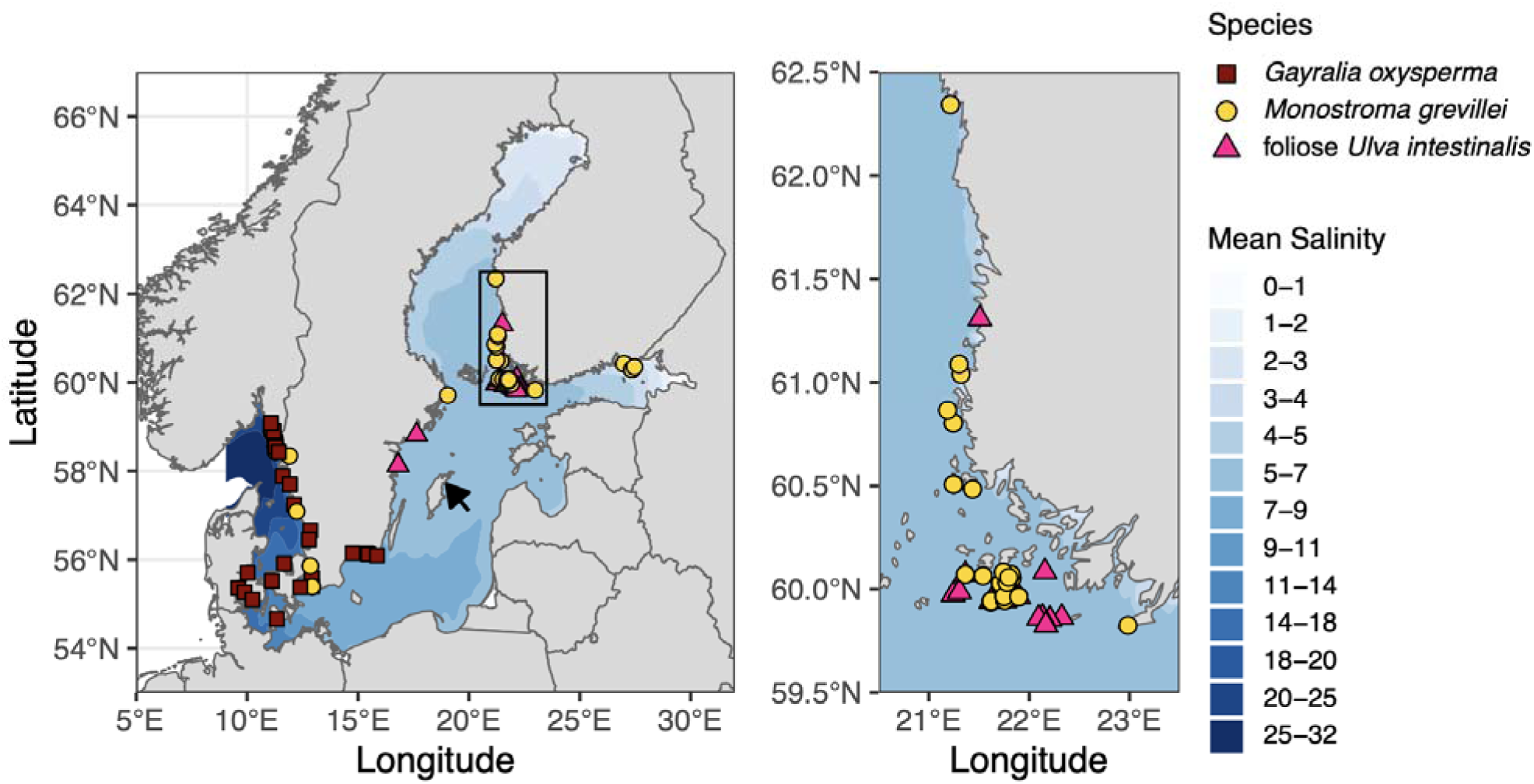
Collection locations and DNA-based identification of foliose, monostromatic green algae in the Baltic Sea. The inset shows an area of dense sampling on the Finnish coast in more detail. The arrow indicates the locality of the material on which the original species description of *Monostroma balticum* was based on.

### DNA extraction, PCR and sequencing of newly collected material

Newly collected silica-dried material was used for DNA extraction following (Steinhagen et al. 2019b) for material from Sweden, Denmark and Norway. DNA extraction for Finnish material was done using the DNeasy Plant Mini Kit (Qiagen, Venlo, The Netherlands) following manufacturer’s instructions. The marker *tuf*A was amplified using PCR in two parts, using newly designed primer combinations (Table 1). PCR was run with the following conditions: initial denaturation 95°C for 4 min, followed by 38 cycles of denaturation 95°C for 1 min, annealing 55°C for 30 s, elongation 72°C for 1 min, followed by a final extension for 7 min. PCR products were cleaned using Exo-SAP (Thermo Scientific, MA, USA) and sanger sequenced by an external provider (Institute for Molecular Medicine Finland FIMM Genomics unit, supported by HiLIFE and Biocenter Finland, University of Helsinki, Finland). Forward and reverse reads were curated and combined to produce full sequences in BioEdit (Hall 1999). Newly produced sequences are published and publicly available in GenBank (submissions SUB16482554 and SUB16485351, GenBank Accession codes being generated by NCBI staff). Alignments used are available at Zenodo (https://doi.org/10.5281/zenodo.22748484).

**Table 1.**
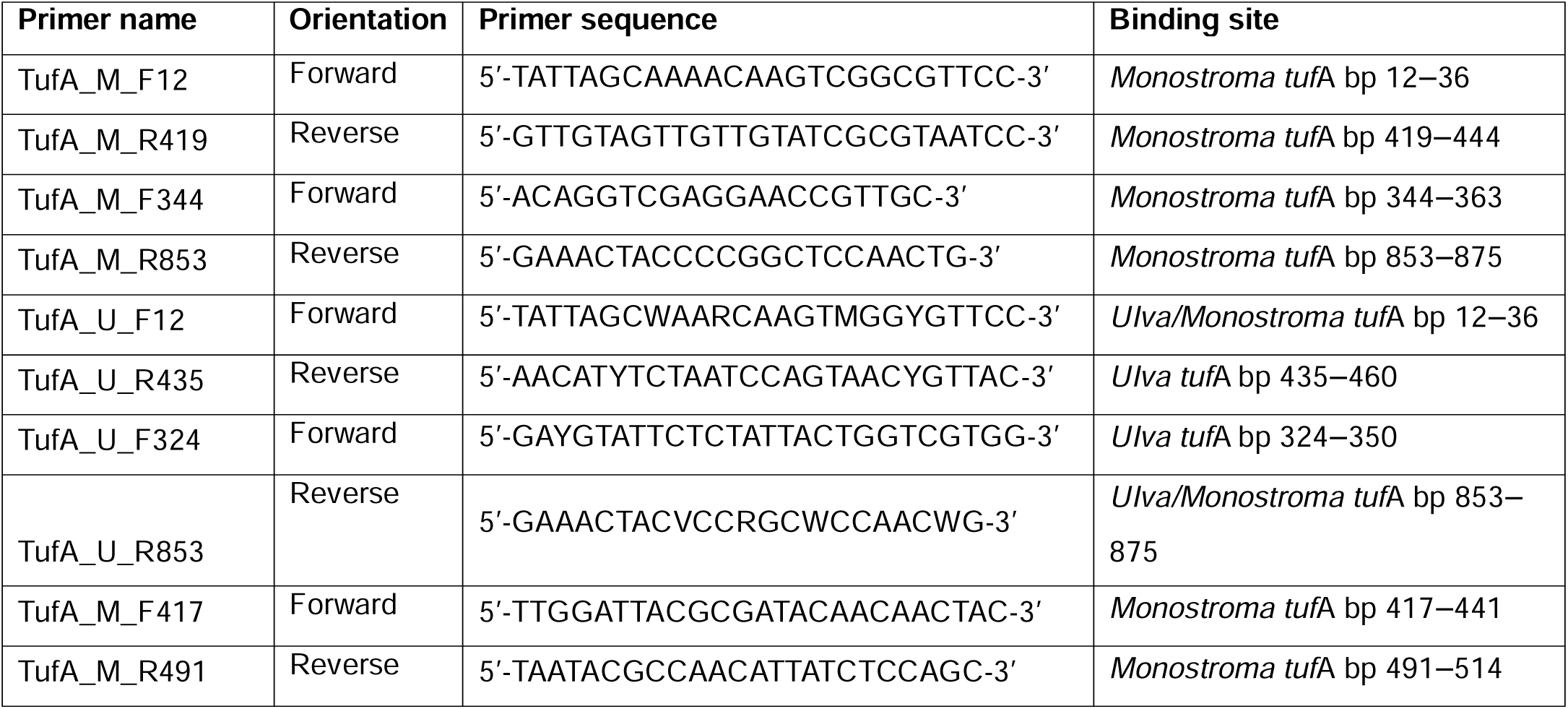
Details of the primers designed and used in this study.

### Sample selection of historical material

*Monostroma balticum* syntype and equivalent material was collected from four specimens in a semi-sterile manner from the Finnish Museum of Natural History herbarium (H) for ancient DNA (aDNA) extraction (Table 2). Samples of c. 1 cm2 dried tissue were carefully selected using gloves and sterile forceps to avoid potential impurities by epiphytes, as well as contamination during sampling. Since the samples were long-term stored as dry herbarium material, no distinct cleaning procedures prior to DNA extraction were conducted, however potential debris (tissue, paper fragments) were removed.

**Table 2.**
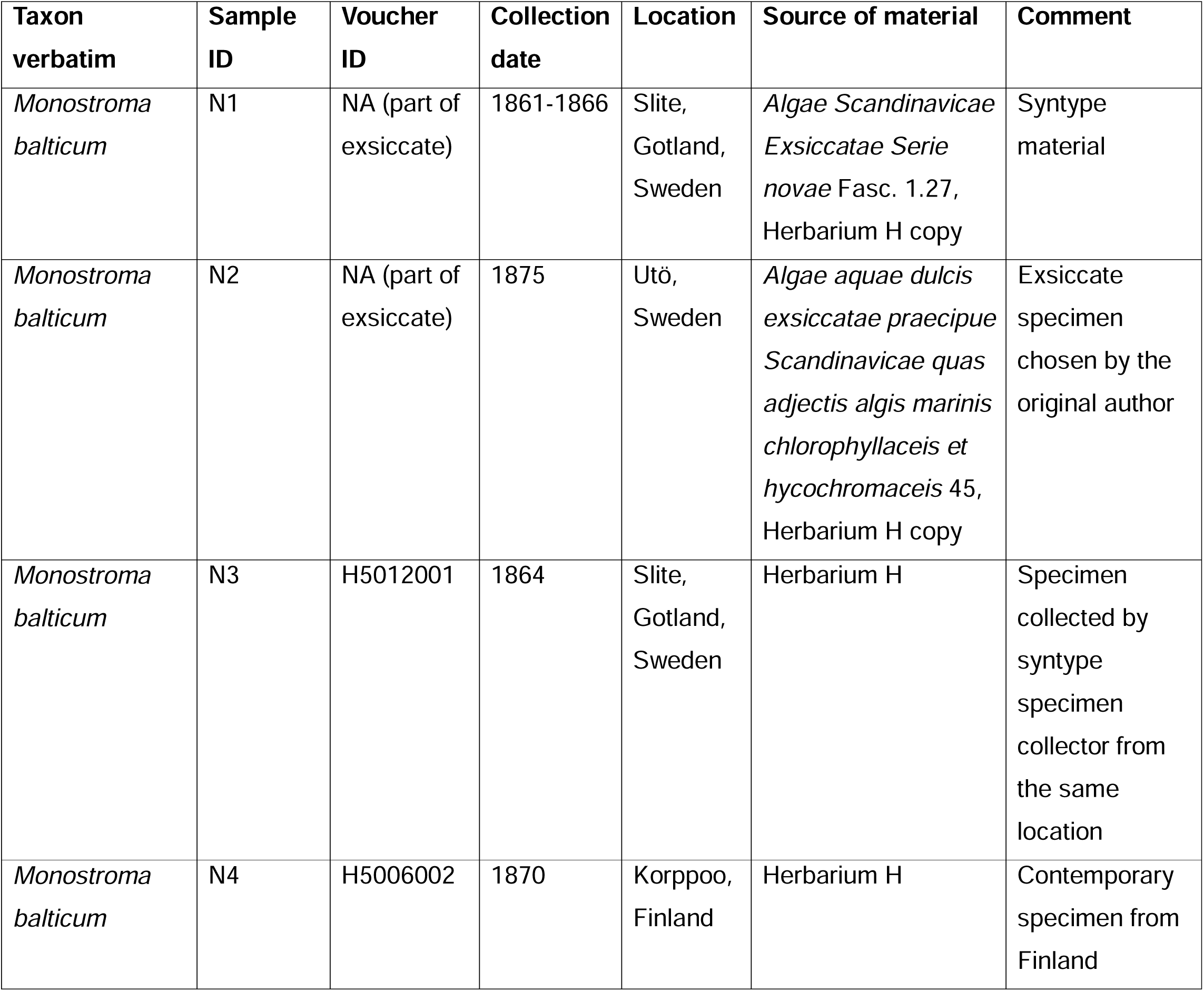
Historical specimens selected for molecular analysis. For photographs of the specimens, see Figs S1-4.

| **Taxon verbatim** | **Sample ID** | **Voucher ID** | **Collection date** | **Location** | **Source of material** | **Comment** |
| --- | --- | --- | --- | --- | --- | --- |
| *Monostroma balticum* | N1 | NA (part of exsiccate) | 1861-1866 | Slite, Gotland, Sweden | *Algae Scandinavicae Exsiccatae Serie novae* Fasc. 1.27, Herbarium H copy | Syntype material |
| *Monostroma balticum* | N2 | NA (part of exsiccate) | 1875 | Utö, Sweden | *Algae aquae dulcis exsiccatae praecipue Scandinavicae quas adjectis algis marinis chlorophyllaceis et hycochromaceis* 45, Herbarium H copy | Exsiccate specimen chosen by the original author |
| *Monostroma balticum* | N3 | H5012001 | 1864 | Slite, Gotland, Sweden | Herbarium H | Specimen collected by syntype specimen collector from the same location |
| *Monostroma balticum* | N4 | H5006002 | 1870 | Korppoo, Finland | Herbarium H | Contemporary specimen from Finland |

### DNA sequencing of historical material

In dry herbarium plant material, chloroplast DNA is prone to fragmentation (McAssey et al. 2023).We therefore applied a museomics approach for sequencing chloroplast DNA from the historic specimens listed in Table 2 focusing on the recovery and sequencing information of short DNA fragments. Ca. 1 cm2 of tissue was used for DNA extraction which followed the guanidine treatment detailed in Straube et al. 2021, which is based on and modified from Sambrook & Russell 2001, as well as Rohland & Hofreiter 2007, Rohland et al. 2010, Dabney et al. 2013, and Basler et al. 2017. Since the tissue samples were derived from dry herbarium material, the PBS buffer pre-extraction was step described in Straube et al. 2021 was skipped. The custom binding apparatus for DNA fragment collection was replaced with preassembled silica spin columns and collection tubes (Rohland et al. 2018). Two negative controls were included in the extraction process flanking the four target samples.

DNA concentrations were measured with a Qubit fluorometer, and additionally maximum peak sizes of the DNA libraries were checked on an Agilent Bioanalyzer using the High Sensitivity DNA Analysis kit. DNA libraries were pooled in equimolar ratios and sequenced on an Illumina MiniSeq instrument using a 75 base pair single-end high-output sequencing kit. Custom sequencing primers were used as described in Paijmans et al. 2017. Extraction and library controls were sequenced along with the samples.

DNA extraction, library preparation, quality checks (qPCR, concentration measurements, and automated electrophoresis) and sequencing of aDNA were performed at the molecular laboratory at the University Museum of Bergen. Laboratory procedures followed recommendations detailed in Fulton and Shapiro 2019 where feasible, including UV-C irradiation, strict decontamination measures, and positive pressure HEPA filtration.

### Bioinformatic processing of historical DNA

Since historical specimens contain generally fragmented DNA, endogenious sequence data yield can be significantly reduced. Following the protocol by Straube et al. 2021, cutadapt v. 1.16 (Martin 2011) was used to trim reads below 30 bp as well as adapter sequences, which allows for the analysis of DNA fragmentation based on calculating average library insert size. Adapters were trimmed with a minimum overlap length between read and adapter of 4 bp and standard bash utilities were applied to investigate average insert size. Untrimmed reads and those with a length of zero were excluded. Trimmed reads were de novo assembled to contigs using the Geneious assembler with medium sensitivity. Both trimmed reads and produced contigs were mapped against reference plastid genome sequences from *Monostroma grevillei* (Johansson et al. 2026, GenBank Accession SUB16485424), *Gayralia oxysperma* (Kützing) K.L.Vinogradova ex Scagel et al. 1989 (Johansson et al. 2026, GenBank Accession SUB16485424) and *Ulva intestinalis* (epitype Historia Muscorum-XXXII, GenBank Accession PX274223, Hughey et al. 2026) using the “map to reference” function in Geneious Prime v2025.0.3 (GraphPad Software LLC, MA, USA). Produced partial *tuf*A contigs were compared to reference sequences and included in the phylogenetic analysis.

### Phylogenetic analysis

Reference sequences from *Ulva* and morphologically similar taxa previously detected from the Baltic Sea were downloaded from GenBank, including all available sequences from type specimens. Additionally, all published *tuf*A sequences representing taxa in *Monostroma* were included. The sequences were trimmed and aligned with MAFFT v.7 with default settings (Katoh et al. 2019). Maximum Likelihood phylogenies were constructed using IQ-TREE2 v.2.4.0 (Minh et al. 2020) using ModelFinder to find the optimal substitution model (Kalyaanamoorthy et al. 2017). Node support was estimated using the ultrafast bootstrap approximation method UFBoot2 (Hoang et al. 2018) with 1000 replicates in IQ-TREE2. The phylogeny was illustrated with TreeViewer (Bianchini & Sánchez-Baracaldo 2024). The phylogenetic analysis was done in two parts: firstly, with representatives of Ulotrichales and Ulvales, and secondly with representatives of Ulvaceae only.

### Microscopy

Fresh material and rehydrated herbarium specimens were studied and photographed with light microscopy (Olympus BX50, Evident Scientific, Tokyo, Japan; Leica flexacam C3, Leica Microsystems, Wetzlar, Germany), mounted in water. Micromophological characters studied included cell and chloroplast shape and size, granules, pyrenoids and cross-sectional features.

## RESULTS

### Molecular identification of newly collected specimens

In total 156 foliose monostromatic specimens were analysed from Finland (n = 91), Sweden (n = 54), Denmark (n = 10) and Norway (n = 1), from which 140 novel sequences were generated (Fig 1, Table S1). The *tuf*A-based phylogenetic analyses were based on two alignments: the Ulotrichales-Ulvales phylogeny (Fig 2) consisted of 32 sequences and 776 nucleotide positions, of which 307 were parsimony informative, and the Ulvaceae phylogeny (Fig 3) consisted of 48 sequences and 673 nucleotide sites, of which 127 were parsimony informative. ModelFinder estimated the best-fit evolutionary model to be TIM3+F+G4 for the Ulotrichales-Ulvales phylogeny and TIM3+F+I+G4 for the Ulvaceae phylogeny.

**Fig 2:**
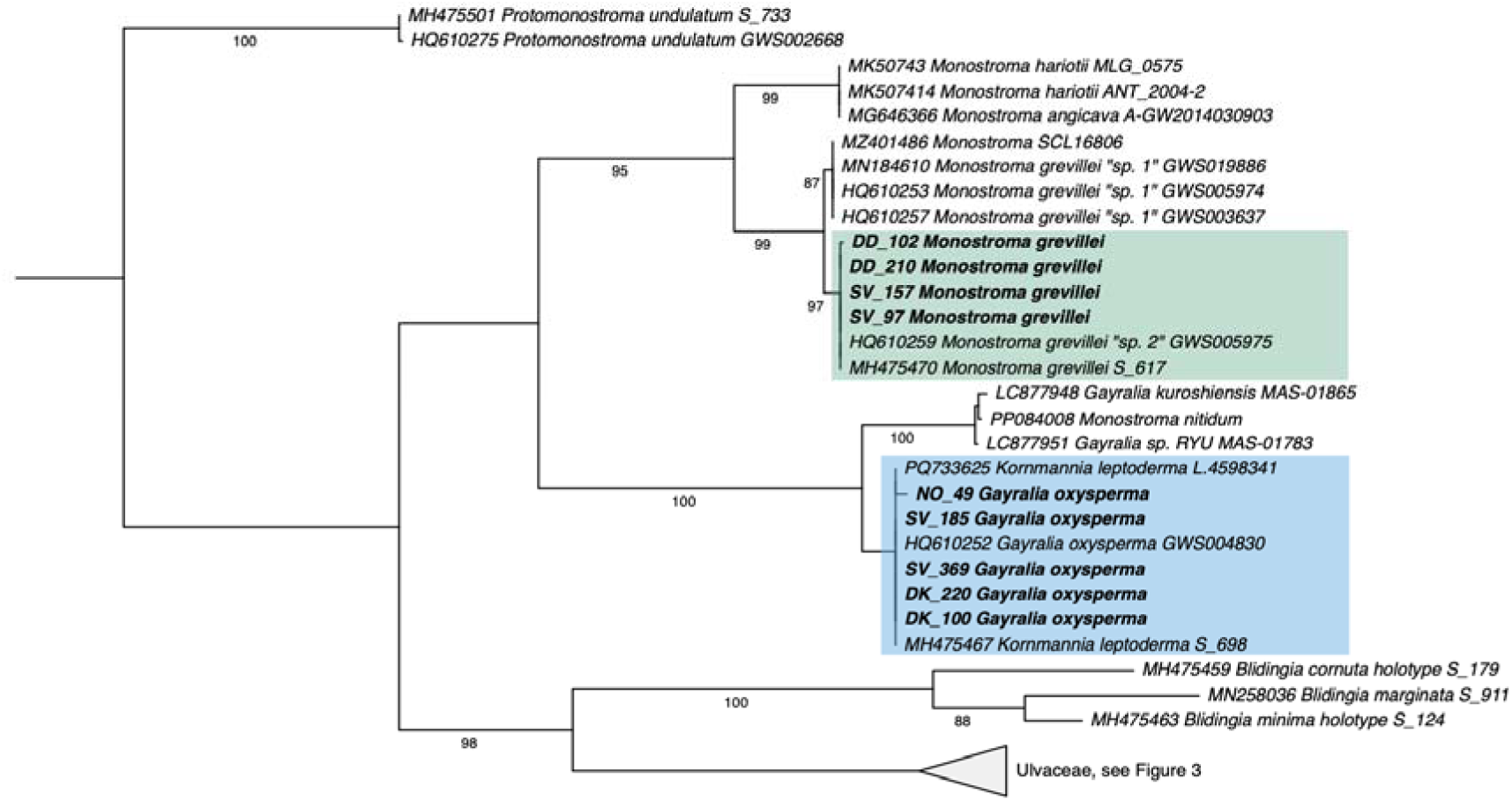
Maximum likelihood phylogeny of selected taxa in Ulotrichales and Ulvales based on the *tuf*A marker gene. Newly generated sequences are shown in bold, up to two per taxon per country when available. Branch values show UFboot2 support values (1000 replicates). Taxa in Ulvaceae is shown in Fig 3.

**Fig 3:**
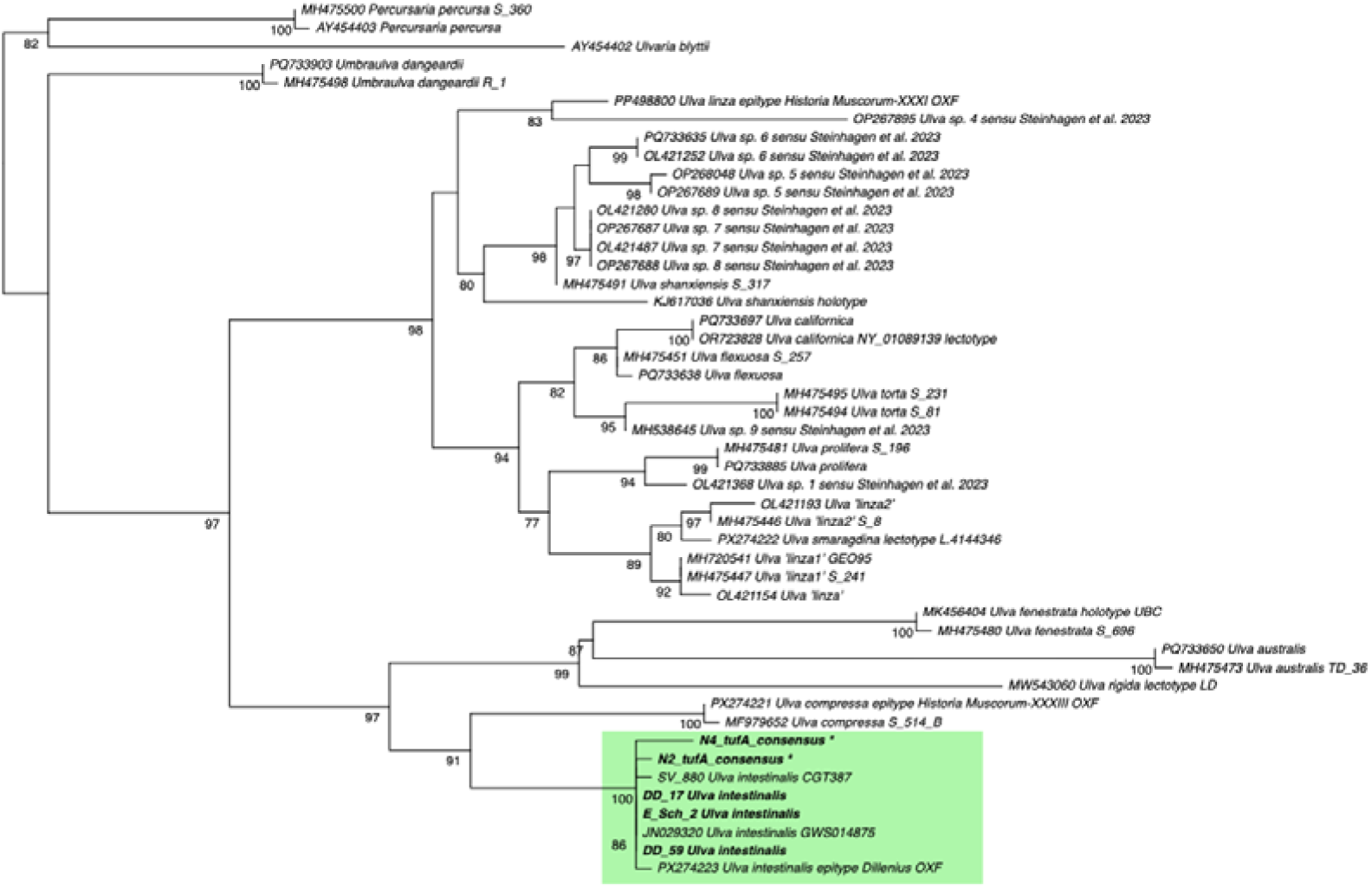
Maximum likelihood phylogeny of selected taxa in Ulvaceae based on the *tuf*A marker gene. Newly generated sequences are shown in bold, up to two per taxon per country when available. Sequences from herbarium specimens are labelled with an asterisk *. Branch values show UFboot2 support values (1000 replicates).

Both phylogenetic analyses place all newly collected specimens to one of three clades: *Ulva intestinalis*, *Gayralia oxysperma* or *Monostroma grevillei* (Figs 2–3). No specimens cluster in a clade within the genus *Monostroma*, which could be considered a separate species *M. balticum*. *U. intestinalis* was recently molecularly verified with sequenced epitype material (Hughey et al. 2026), confirming that the clade labelled *U. intestinalis* in our phylogeny represents this taxonomically accepted entity. In *M. grevillei,* two clades are recovered, “sp. 1” and “sp. 2” sensu Saunders & Kucera 2010, our sequences fall into the more cosmopolitan “sp. 2”. The correct name for the *Gayralia oxysperma* clade is unclear, as this clade has been referred to as *Kornmannia leptoderma* in previous publications (Steinhagen et al. 2019b, Weinberger et al. 2019), however, recent results and the phylogenetic placement in Ulotrichales suggest the clade should be called *G. oxysperma* (van der Loos et al. 2025). For labelling these two clades, we use the species names *Monostroma grevillei* and *Gayralia oxysperma* tentatively, until confirmatory reference sequences from type material becomes available.

Foliose *Ulva intestinalis* is restricted to the inner Baltic Sea basin in our dataset, most specimens collected from the Archipelago Sea and the Eastern Swedish coast. All foliose *U. intestinalis* collections are made in low salinity conditions (<7). This species is observed in the summer season (June-September) as well as the spring season (April). *Gayralia oxysperma* is restricted to the southern Baltic Sea, the Danish straits, Kattegat and Skagerrak to salinities generally >7, and found throughout the season (January-August). *Monostroma grevillei* is found across the sampling region, from the oceanic conditions in the Atlantic-Baltic Sea transition area to low salinity conditions in the Inner Baltic Sea. The easternmost specimens are found at salinity 4 in the Gulf of Finland, whereas the northern range edge of the continuous population at c. 61°N lies in salinity of c. 5. The most northern specimens at 62°N are an isolated population located next to a fish farm, and despite extensive searches, no individuals have been found between the continuous range and this isolated population in the Finnish west coast. The species is observed in the spring season, latest specimens collected in early May from the Gulf of Finland.

### *Monostroma balticum* Wittrock matches specimens of foliose *Ulva intestinalis*, and not *Monostroma grevillei* or *Gayralia oxysperma*

In the original description by Wittrock 1866, the thallus is described as membrane-like, irregular in shape, wrinkled and creased, pierced with holes of various shapes and sizes. The blade of the thallus is described as thin (28–33 µm) but relatively stiff and whitish green. Thallus size is described to be 3–4 tum long and 2–3 tum wide. Tum is an old Swedish measure of length, the absolute length of which has varied across years and locations in Sweden. According to Engström 1883, a tum during the years 1855–1880 (Wittrock’s dissertation was published 1866) corresponded to 2.97 cm, making the measurements of the thallus approximately 9–12 cm long and 6–9 cm wide. Wittrock notes that the species is known only detached, drifting freely in water (but argues, that it can most likely be found attached in juvenile form). No holdfast structure is described, and the thalli are thought to be completely homogeneous throughout the blade. Further workers describe the macromorphology in similar terms to Wittrock, emphasising the fact that the species is found only free-floating – making this feature often the diagnostic feature for species identification (Svedelius 1901, Levring 1940, Wærn 1952, Ravanko 1968, Tolstoy & Österlund 2003). Notably, the size of the thallus has been cited to be larger than Wittrock’s original publication. Holes or fenestrae are mentioned only in the original Wittrock 1866 description and not by any later workers. Macromorphologically, newly collected specimens which cluster with *Ulva intestinalis* resemble the original Wittrock description well (Figs 4A-E).

**Fig 4.**
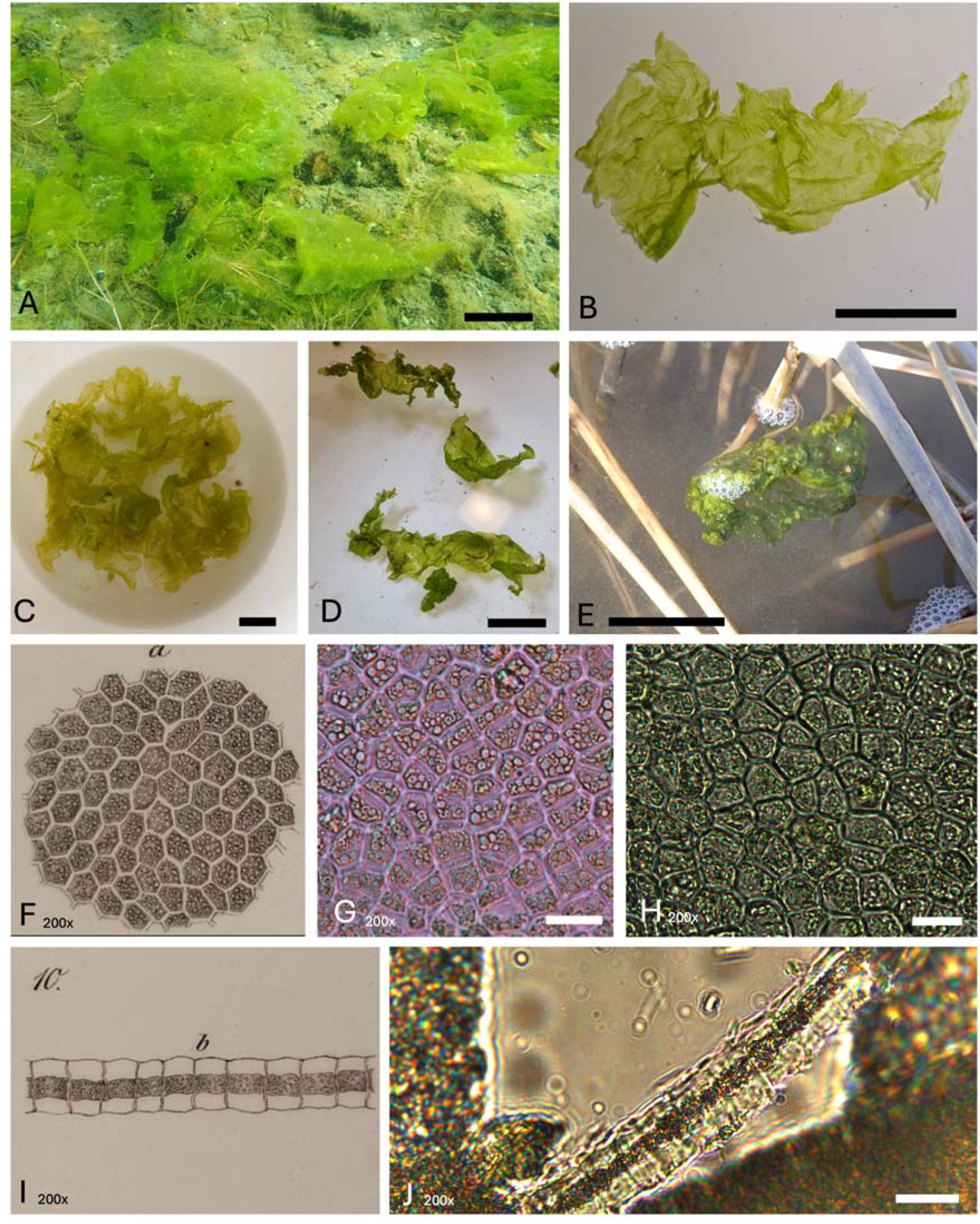
Morphological features of foliose, monostromatic green algae molecularly identified as *Ulva intestinalis*. A-C) typical free-floating late season specimens. D-E) dark green early season specimens, collected among ice. F) original illustration of the cellular structure in *M. balticum* from Wittrock 1866. G-H) cellular structure of fresh material. I) original illustration of the cross-section of the blade from Wittrock 1866. J) cross-section of fresh material. Scale bars 4A,C,E, 5 cm; 4B,D; 2.5 cm; 4G-J, 20 μm.

Considering micromorphology, Wittrock describes the cells to be 5–7-cornered, often hexagonal, fully green, tightly packed and without order. The chloroplast is described plate-like, and in cross section it forms a distinct 8–11 µm band-like structure throughout the cell’s width. From the top view cells are 15–21 µm long and 9–15 µm wide, and in cross-section 27.5–32.5 µm high. Wittrock mentions multiple starch granules in each cell. Detailed drawings were produced which illustrate the cellular structure from top view and cross-section (Figs 4F, I). Further workers generally agree with Wittrock’s original description, whereas the individual measurements vary slightly when cited. In general, the descriptions of cellular structure are vaguer in works after Wittrock. Starch granules or pyrenoids are not mentioned after Wittrock, even if granular structures are clearly visible in his drawings (Fig 4F). Few workers mention a thickening of the outer cellular wall (Svedelius 1901, Levring 1940), while in others the cell walls are described as thin (Tolstoy & Österlund 2003). Cross-sectional features are not used in contemporary sources.

While the description of micromorphology of the taxon is vague, especially in contemporary treatments, the micromorphology of newly collected specimens clustering with *Ulva intestinalis* agrees well with the original description (Fig 4G-H, J). Blade cross-section structure with a thin, green band consisting of the chloroplast, considered diagnostic in Wittrock’s treatment, can be detected in fresh specimens (Fig 4J).

The ecology and phenology of foliose *Ulva intestinalis* generally match to the original Wittrock description. However, some specimens from Finland were collected in April, outside of the known *Monostroma balticum* occurrence phenology of late summer. These specimens show a thicker and denser, darker green blade, but otherwise fit the description of *M. balticum* (Figs 4D-E). These specimens most likely represent overwintered foliose *U. intestinalis* thalli.

Specimens clustering with *Monostroma grevillei* morphologically match the description of this well-studied cosmopolitan species (Burrows 1991, Nielsen et al. 2022). The majority of *M. grevillei* specimens are collected as attached, while some specimens are free-floating. These unattached specimens are typically relatively large, thin, fragmented and co-occur with attached *M. grevillei* in late spring. We consider these large, mature individuals of *M. grevillei* which have become free-floating e.g. via physical disruption by wave action and not the taxon originally described as *Monostroma balticum* by Wittrock. Wittrock was very familiar with *M. grevillei* and discusses this species in length (Wittrock 1866). Thus, we consider it unlikely that a mature, unattached thallus of this species would have been mistakenly described as a novel species in his treatment (Wittrock 1866).

Specimens clustering with *Gayralia oxysperma* follow the morphology described in (Steinhagen et al. 2019b, Weinberger et al. 2019). These specimens occur almost exclusively attached, with a single case of an unattached specimen collected, co-occurring with attached forms. Micromorphological features of *G. oxysperma* do not correspond well to the Wittrock description of *M. balticum*, especially as *G. oxysperma* has 1(–2) central pyrenoids and a more rounded cell shape. Wittrock discusses this species as well in length, as *Monostroma oxycoccum* (Wittrock 1886). The range of *G. oxysperma* detected here or reported previously does not reach the locations where the original *M. balticum* material studied by Wittrock was collected, or where *M. balticum* is typically reported from.

Based on the combination of macro- and micromorphological, ecological, phenological and distributional data, we conclude that of the three molecularly characterised taxa detected in our sampling, the foliose *Ulva intestinalis* best matches the original Wittrock description of *Monostroma balticum*.

### Historical *Monostroma balticum* material was molecularly identified as *Ulva intestinalis*

Two out of the four historical specimens analysed yielded enough reads for a molecular identification. Sequencing of specimen N2 resulted in 1 334 378 raw reads (average insert size 41.2 bp), 166 373 trimmed reads (assembled to 6637 contigs), of which 4617 were mapped to the *Ulva intestinalis* epitype chloroplast genome (PX274223, Hughey et al. 2026). This produced a full-length *tuf*A sequence apart from gaps of 8 bp and 12 bp and 12 ambiguous bases (GenBank Accession SUB16485438). Sequencing of specimen N4 resulted in 1 773 405 raw reads (average insert size 45.1 bp), 131 765 trimmed reads (assembled to 9907 contigs) of which 3169 were mapped to the same *U. intestinalis* epitype. This produced a partial full-length *tuf*A sequence, which included 6 gap regions (lengths 3-37 bp) and 15 ambiguous bases (GenBank accession SUB16485438). Similar mappings to reference genomes of *Monostroma grevillei* and *Gayralia oxysperma* (Johansson et al. 2026) produced poorer or no *tuf*A sequences, which have the same sequence as when mapped to the *Ulva* genome. Specimens N1 and N3 yielded 3099 and 50 867 trimmed reads, respectively, which did not map well to any reference genomes used, and produced no *tuf*A contigs. Reads from these samples were primarily of bacterial origin.

While Wittrock 1866 does not designate a holotype specimen, the reference to an exsiccate gathering can be considered a designation of a syntype series. We have confirmed that exsiccate specimens from H (N1, Fig S1), PC and L form a visibly homogeneous collection with shared collection data and labels, thus the entire exsiccate gathering can be considered a syntype series under Art. 9.6. (Turland et al. 2025). The exsiccate material originates from the port of Slite, Gotland, Sweden (Wittrock 1866).

While the available reads of the syntype material (N1, Table 2, Fig S1) did not allow for molecular identification, the specimen N2 from a later exsiccate series prepared by Wittrock and co-authors is molecularly confirmed to be a foliose specimen of *Ulva intestinalis* (Fig 3, Fig S2). This specimen originates from a strait between Rånö and Ålö on Utö island, Sweden, ca. 200 km from the syntype location. The gathering of this exsiccate series was made by the same collector as the syntype series (T. O. B. N. Krok) and given that Wittrock himself authored this later exsiccate series, it is likely that these specimens belong to the same species concept of *Monostroma balticum* Wittrock. The specimen N4 (Fig S4), collected at around the same time from the Finnish Archipelago Sea by Fredrik Elfving, is similarly molecularly confirmed to be a foliose form of *U. intestinalis* (Fig 3). We consider this evidence to be enough to treat *M. balticum* as a monostromatic, free-floating foliose growth form of *U. intestinalis*, and thus designate *M. balticum* as a heterotypic synonym of *U. intestinalis*:

(This is a preprint, and we do not consider nomenclatural acts in this publication effectively final under Art. 30.2. (Turland et al. 2025))

### TAXONOMY

#### *Ulva intestinalis* Linnaeus 1753: 1163

EPITYPE: OXF, Tremella marina tubulosa, Woolwich, London, England, collected between 1721 and 1741, no habitat data, leg. Johann Jakob Dillenius (designated by Hughey et al. 2026)

LECTOTYPE: Dillenius, 1742: 47, plate 9, figure 7 (designated by Blomster et al. 1998).

### HETEROTYPIC SYNONYMS

#### *Monostroma balticum* Wittrock 1866: 48–49

TYPE/SYNTYPES: Slite port, Gotland, Sweden, July, collected between 1861 and 1866, T. O. Krok in Alg. Scand. Exs. 1: 27 (H, PC, L…). See Fig S1 for a photograph of the type.

#### Phycoseris planifolia Kützing 1843: 297

HOLOTYPE: L: L.4144341, Timavo River, Malfacone, Italy, 1835, no habitat data, collector unknown (Hughey et al. 2026).

These are the only proposed heterotypic synonyms of *U. intestinalis* which have their type specimens or equivalent material sequenced (Hughey et al. 2026), while AlgaeBase currently recognises 11 homotypic and 14 heterotypic synonyms (Guiry & Guiry 2026).

## DISCUSSION

Our findings are a clear example that species of green macroalgae in *Ulva* and morphologically similar genera can be phenotypically highly variable. There is increasing recognition that macroalgal phenotypic plasticity can be extensive. However, typically reported phenotype shifts are continuous, such as changes in thallus size and branching patterns (Coleman & Martone 2024 and references therein), pneumatocysts (Stewart 2006) or changes in reproductive features (Johansson et al. 2017). Plasticity of entire growth forms are much more rarely reported (see. e.g. Monro & Poore 2009, Pongparadon et al. 2020)

*Ulva intestinalis* grows typically in a tubular form in the Baltic Sea (Nielsen et al. 2022, Steinhagen et al. 2023). Previous reports of foliose *U. intestinalis* do exist from the Baltic Sea, especially in the context of green tides (Bäck et al. 2000, Blomster et al. 2002, Steinhagen et al. 2019b). These reports clearly demonstrate an aberrant, monostromatic growth form or morphotype of molecularly confirmed *Ulva (Enteromorpha) intestinalis*, with a similar morphology and ecology to the *Monostroma balticum* original description. The conspecificity of these *Ulva* green tides to *M. balticum* was raised speculatively (e.g. (Blomster et al. 2002), but until now, molecular data from *M. balticum* has been lacking to confirm this speculation. Based on our results, it now seems likely that monostromatic green tides in the inner Baltic Sea are mainly caused by foliose *U. intestinalis*, and that previous reports of green tides attributed to *M. balticum* most likely represent *U. intestinalis* (Mathiesen & Mathiesen 1992). Despite extensive searches, we have not found any reports of a foliose morphotype of *U. intestinalis* outside of the Baltic Sea.

Other *Ulva* species with a primary tubular attached growth form are known to show an occasional secondary foliose growth form. These include, for instance, *U. smaragdina* (Kützing) Hughey, Maggs, L.M.Loos, S.A.Harris & P.W.Gabrielson (previously often reported as *U. linza*, Hughey et al. 2026) and *U. compressa* Linnaeus (Tan et al. 1999, Hofmann et al. 2010, Steinhagen et al. 2019c, 2023). However, these foliose blades are typically distromatic, unlike the monostromatic growth form of *U. intestinalis* detected in our study (Fort et al. 2021).

The foliose morphotype of *Ulva intestinalis* seems to be linked to the low salinity conditions of the inner Baltic Sea, whereas the tubular form is seen ubiquitously, across a wide salinity gradient, and is always co-occurring with the foliose morphotype. Similarly, the foliose form of *U. compressa* in Atlantic USA is associated with lower salinity estuarine areas, while the typical tubular form is more oceanic (Hofmann et al. 2010). In contrast, across species of *Ulva*, foliose blade-forming species are usually associated with high salinities, whereas low salinities are predominantly inhabited by tubular representatives (Rybak 2018). This suggests that salinity may drive the phenotype of *Ulva* in different ways on different scales: while a foliose growth form may be generally favoured in high-salinity conditions (Rybak 2018), within a predominantly tubular taxon, low-salinity conditions may trigger a specific low-salinity foliose morphotype. Low salinity, or fluctuations in salinity, are linked to other phenotypic changes in *U. intestinalis*, such as changes in branching pattern within the tubular form (Reed & Russell 1978).

This raises speculation that low salinity, perhaps together with some other hitherto unknown environmental factors or conditions, causes a growth form shift from a tubular source population to a foliose accumulation in *Ulva intestinalis*, at least in the Baltic. Growth experiments by Blomster et al. (2002) demonstrate that the foliose growth form may give rise to a tubular thallus in laboratory conditions, but the opposite was not achieved in their experiments. Potentially some large *U. intestinalis* individuals, when becoming unattached and unravelled in specific environmental conditions, can give rise to foliose sheets. Growth experiments by Orvokki Ravanko (1969), claiming that *Monostroma balticum* from the Finnish archipelago correspond to the juvenile ontogeny of *M. grevillei* most likely represent actual *M. grevillei* and not foliose *U. intestinalis*.

Given the large size of some foliose specimens, it is likely that foliose forms keep growing vegetatively in free-floating form. This is also evident on their morphology, as the monostromatic foliose sheets are not planar, but irregularly undulating – showing evidence of thallus growth. Additionally, dividing cells are frequently observed by microscopy. We have not observed any signs of sexual reproduction in foliose *U. intestinalis* using microscopy, while sporulation is frequent in even small, attached thalli of the tubular growth form. It is unknown if the foliose growth from is a gametophyte or a sporophyte, or if both isomorphic generations can form foliose morphotypes. Further investigations of the foliose morphotype should be conducted, including growth experiments, which could detect what factors, be it environmental, physiological or genetical, give rise to the tubular-to-foliose growth from transition.

Population genetic analyses will be important to further elucidate the origin and persistence of the foliose morphotype. Comparing the genetic diversity and population structure of co-occurring tubular and foliose individuals could reveal whether foliose thalli represent genetically differentiated populations or whether the two morphotypes belong to a single, interbreeding population. Such analyses could also help determine whether large free-floating foliose thalli originate from repeated independent morphological transitions or represent the vegetative proliferation of a limited number of genotypes. In combination with controlled growth experiments, population genomic approaches could therefore help disentangle the relative contributions of genetic differentiation, environmental conditions, and phenotypic plasticity to the formation of the foliose morphotype.

The foliose forms can be long-living, as some specimens collected are overwintered – this was shown also by Bäck et al. 2000 and Blomster et al. 2002. We consider it unlikely that these overwintering thalli are perennial. It seems likely that the foliose morphotype forms more-or-less transient and sporadic populations, and do not form a distinct sub-population in the larger *U. intestinalis* population. However, in e.g. *Fucus*, asexual, vegetatively reproducing free-drifting sub-populations have been demonstrated to be genetically distinct populations with a distinct genetic structure, comparing to the sexually reproducing main population (see e.g. Preston et al. 2025 and references therein).

Based on current data, this special morphotype seems restricted to the Baltic Sea. This fits the claim that was made for *Monostroma balticum*, that it was a local endemic of the region. Further work should explore if similar foliose *Ulva intestinalis* forms could be found in other areas with similar salinity gradients. The ecological role of foliose *U. intestinalis* in the Baltic Sea is unclear. As a potential green tide species it could be of concern in certain situations, but in other situations it may play specific role in the Baltic Sea ecosystem that is separate to the more common tubular form. For instance, in winter and spring conditions, it may be an important food web component when other green macroalgae are sparsely available.

Our results clearly demonstrate that morphological features alone are not reliable in species identification of cryptic green macroalgae. Especially in cases of aberrant or unusual morphotypes, identification keys based on “standard” growth form morphologies are typically misleading at best and unusable at worst. In these cases, DNA-based approaches appear as the best way to make reliable identifications. Increasing efforts at characterising historical species concepts molecularly, and linking them to contemporary sequences and specimes, are paving the way towards a more rigorous understanding of algal biodiversity in the Baltic Sea and beyond.

## ACKNOWLEDGEMENTS

We thank the staff and field personnel from the Metsähallitus Parks & Wildlife Finland and the Finnish Inventory Programme for Underwater Marine Diversity (VELMU), especially Kevin O’Brien, Anna Lyssenko, Ari Laine, Erika von Essen, Sara Karvo, Maiju Lanki, Essi Keskinen, Paula Ruokolainen and Joonas Hoikkala. We thank Kyung Min Lee from the Finnish Museum of Natural History Luomus for coordinating the laboratory work. We also thank Marja Koistinen, Jaana Haapala and Henry Väre from the Finnish Museum of Natural History Luomus Botany & Mycology unit for access to herbarium specimens, and Alexander Sennikov & Teuvo Ahti for advice on nomenclatural issues. We thank Ellen Schagerström and Lena Kautsky for their support with specimen provision and collection. Additionally, we thank Erika von Essen, Paula Ruokolainen and Ellen Shcagerström for photographs in Figure 4. We would like to express our sincere thanks to Louise Maria Lindblom (University Museum of Bergen, Norway) for support during museomics labwork and Mayre Ellen Rodrigues de Sousa (University of São Paulo) for help with initial bioinformatics. The authors wish to acknowledge CSC – IT Center for Science, Finland, for computational resources.

## CREDIT ROLES

Niko R. Johansson: conceptualization (equal); field sampling (supporting); laboratory work (supporting); analysis (lead); writing – original draft preparation (lead); writing – review & editing (lead). Jaanika Blomster: conceptualization (equal); supervision (equal); writing – review & editing (equal). Tytti Wärri: field sampling (equal); writing – review & editing (supporting). Elina Laiho: laboratory work (equal); primer design (lead); writing – review & editing (supporting). Elena Schrofner-Brunner: laboratory work (supporting); writing – review & editing (supporting). Nicolas Straube: laboratory work (equal); writing – review & editing (supporting). Cessa Rauch: laboratory work (supporting); writing – review & editing (supporting). Sophie Steinhagen: conceptualization (equal); field sampling (equal); laboratory work (equal); supervision (equal); writing – review & editing (equal).

## DECLARATION OF GENERATIVE AI USE

The authors report generative AI was not used in their research or preparation of this manuscript.

## DATA AVAILABILITY

All produced sequence data is uploaded and freely accessible in GenBank (SUB16482554 and SUB16485351, GenBank Accession codes TBC upon designation by NCBI staff). Alignments used are deposited in Zenodo: https://doi.org/10.5281/zenodo.22748484. All voucher specimens are stored in public herbaria (see Table S1). Data available from authors upon request before database processing is complete.

