## Supplementary Figures for "The proposed Baltic Sea endemic green alga *Monostroma balticum* represents a foliose morphotype of *Ulva intestinalis* (Ulvales, Ulvophyceae)"

Figure S1. N1**:** the Helsinki copy of the *Algae Scandinavicae Exsiccatae Serie novae* exsiccate Fasc. 1.27, a specimen labelled *Ulva lactuca Ag. Var.?* (syntype material from Slite, Gotland, Sweden by T. O. Krok).

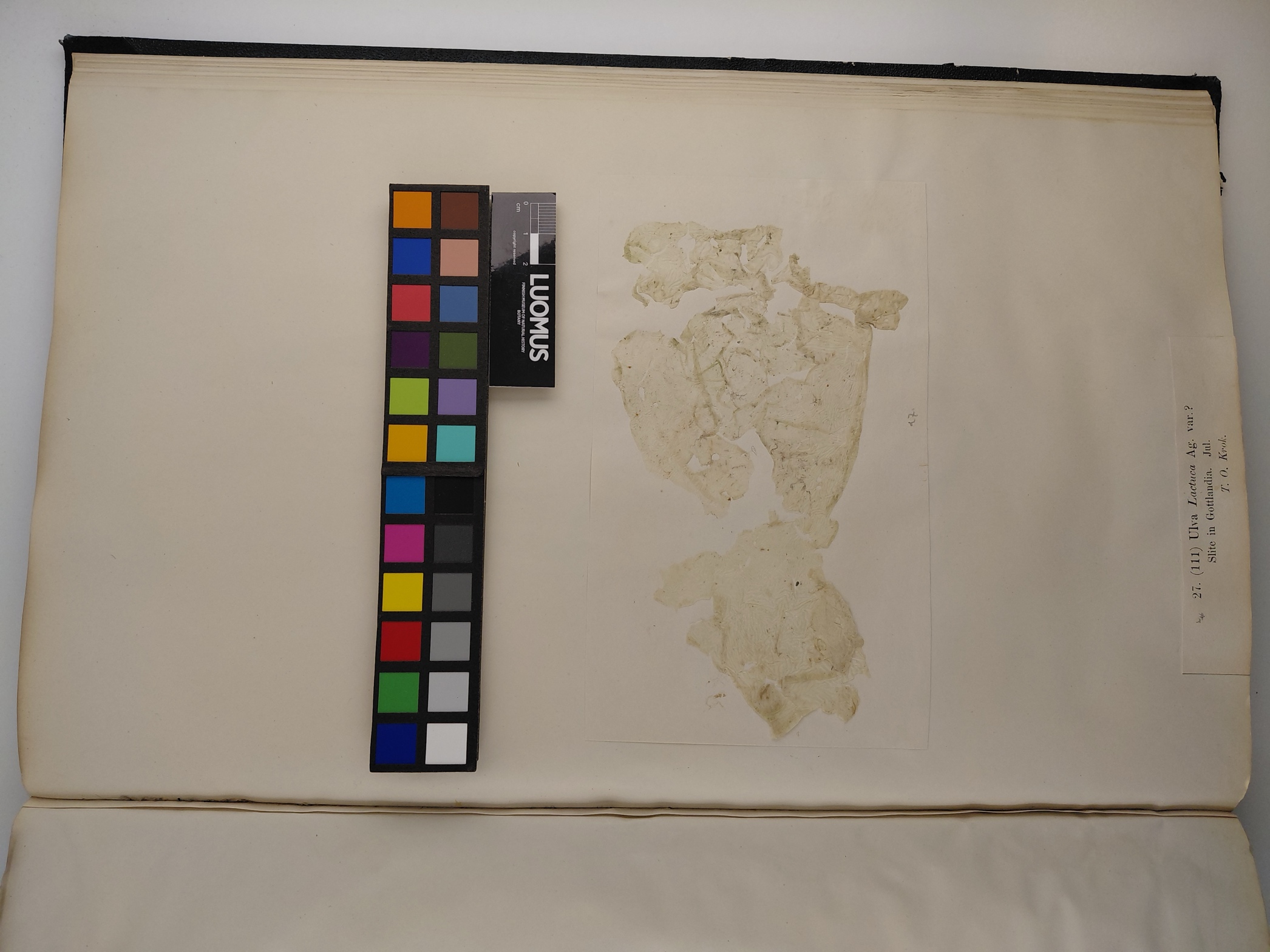

Figure S2. N2: the Helsinki copy of the exsiccate *Algae aquae dulcis exsiccatae praecipue Scandinavicae quas adjectis algis marinis chlorophyllaceis et hycochromaceis* by Wittrock and co-authors, specimen 45 labelled *Monostroma balticum,* material from Utö, Sweden by T. O. B. N. Krok, 1875.

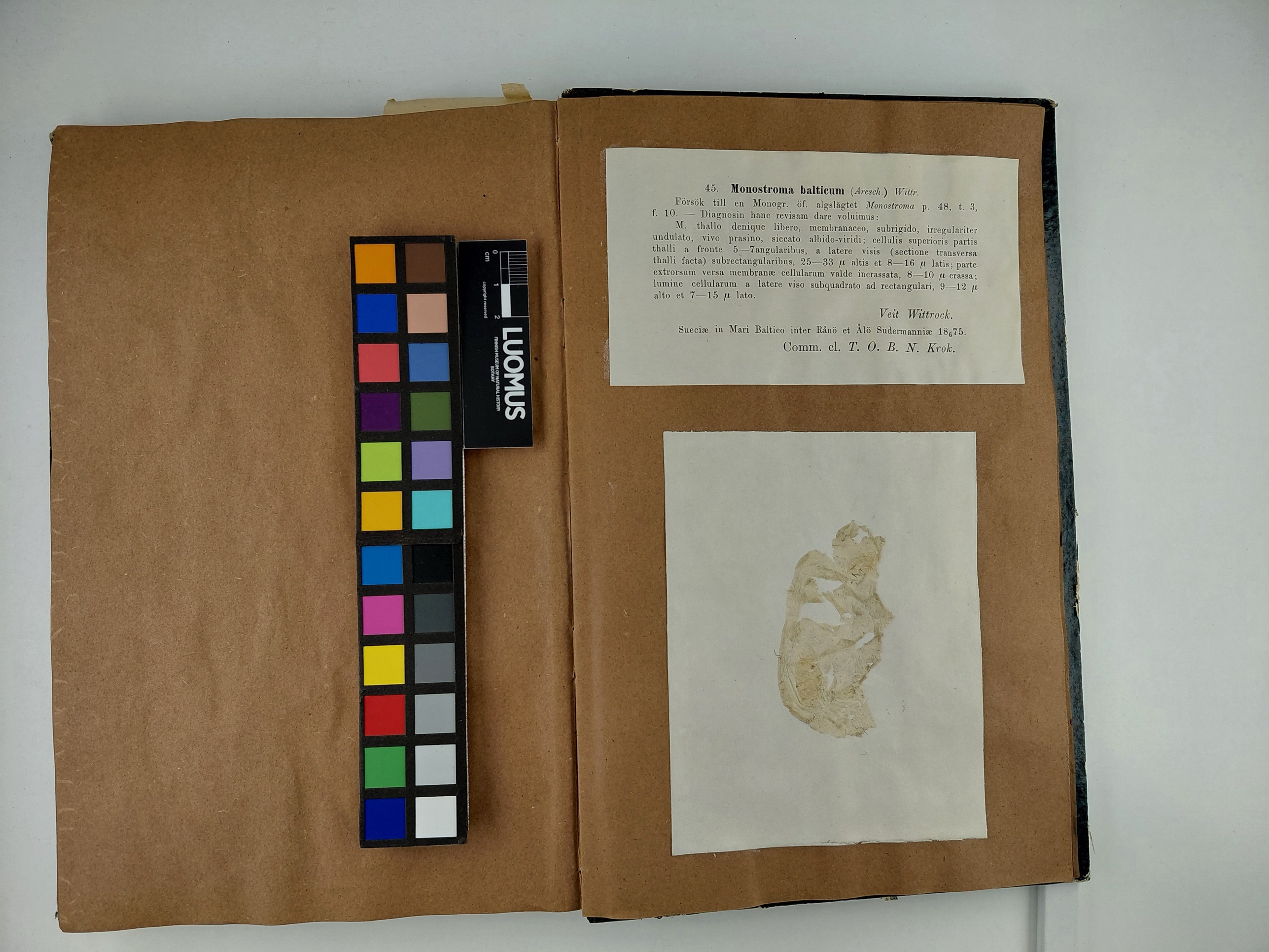

Figure S3. N3: specimen H5012001 from Slite, Gotland by P. T. Cleve, 1864, collected before Wittrock’s original description from the type location.

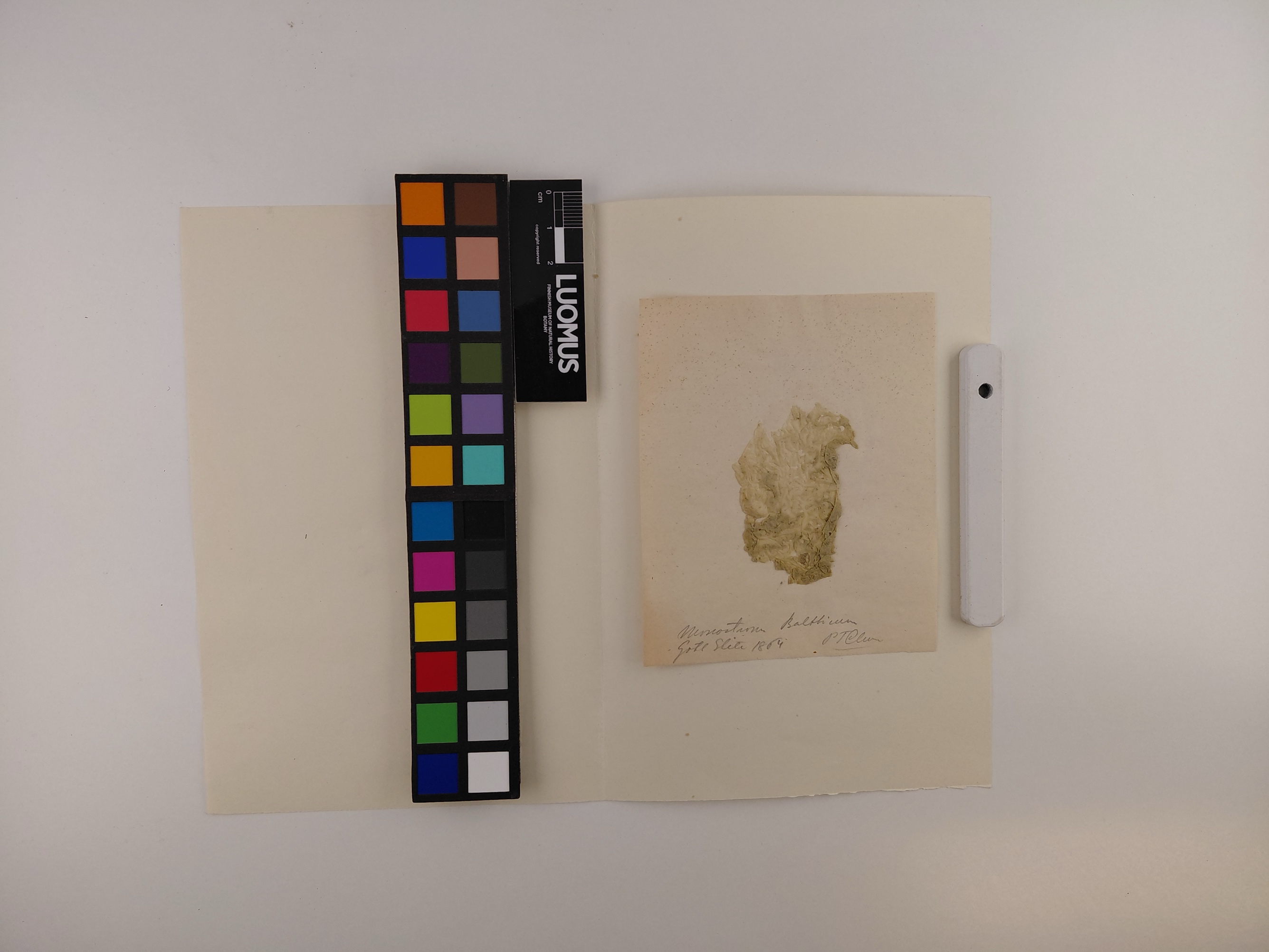

Figure S4. N4: specimen H5006002, a 1870 sample from the Finnish coast (Långvik, Korppoo) by Fredrik Elfving.

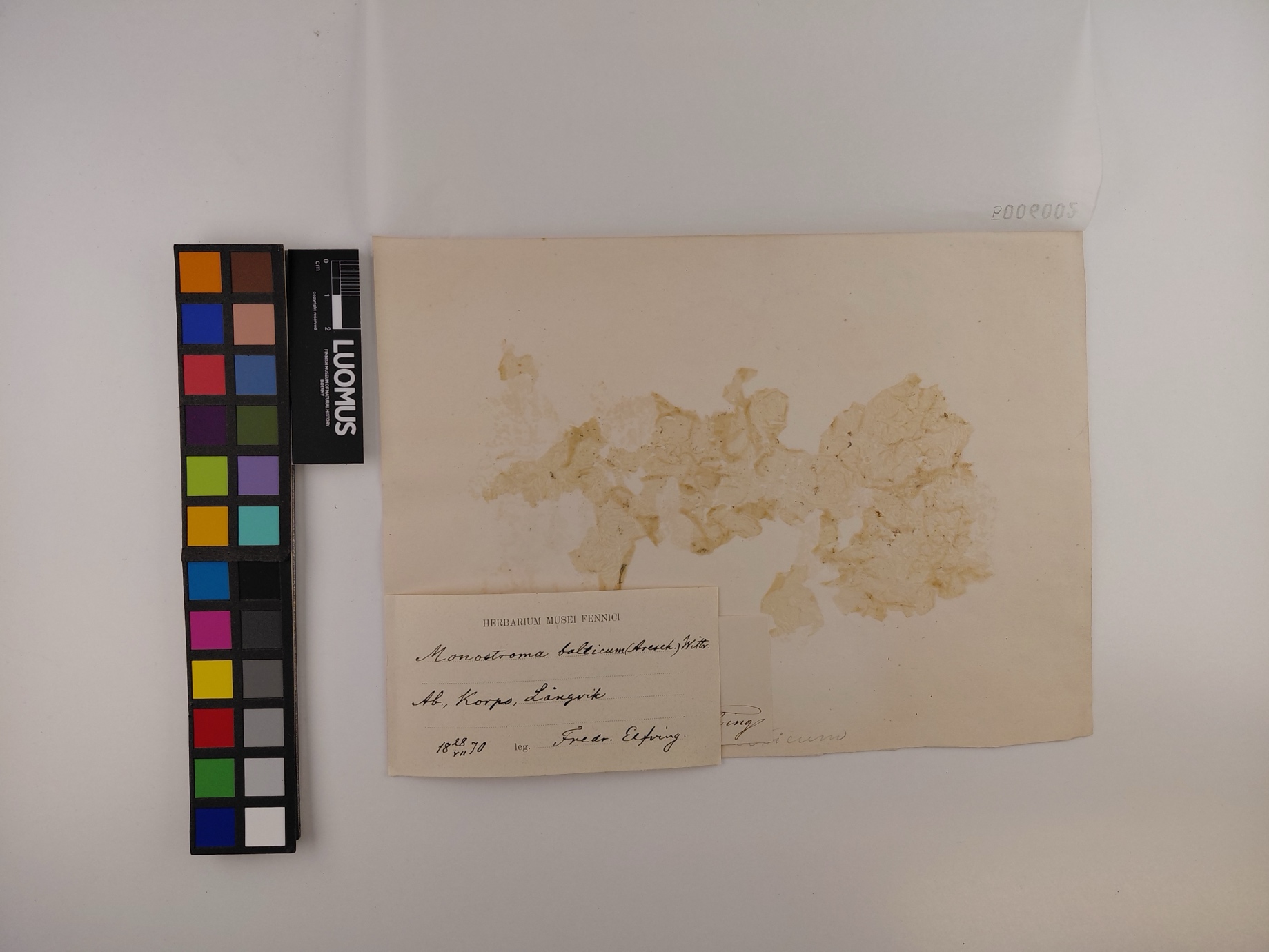
